# SARS-CoV-2 ORF8 modulates lung inflammation and clinical disease progression

**DOI:** 10.1101/2023.09.08.556788

**Authors:** Marisa E. McGrath, Yong Xue, Louis Taylor, Carly Dillen, Jeremy Ardanuy, Norberto Gonzalez-Juarbe, Lauren Baracco, Raymond Kim, Rebecca Hart, Nacyra Assad-Garcia, Sanjay Vashee, Matthew B. Frieman

## Abstract

The virus severe acute respiratory syndrome coronavirus 2, SARS-CoV-2, is the causative agent of the current COVID-19 pandemic. It possesses a large 30 kilobase (kb) genome that encodes structural, non-structural, and accessory proteins. Although not necessary to cause disease, these accessory proteins are known to influence viral replication and pathogenesis. Through the synthesis of novel infectious clones of SARS-CoV-2 that lack one or more of the accessory proteins of the virus, we have found that one of these accessory proteins, ORF8, is critical for the modulation of the host inflammatory response. Mice infected with a SARS-CoV-2 virus lacking ORF8 exhibit increased weight loss and exacerbated macrophage infiltration into the lungs. Additionally, infection of mice with recombinant SARS-CoV-2 viruses encoding ORF8 mutations found in variants of concern reveal that naturally occurring mutations in this protein influence disease severity. Our studies with a virus lacking this ORF8 protein and viruses possessing naturally occurring point mutations in this protein demonstrate that this protein impacts pathogenesis.

**Significance:** Since its emergence in 2019, SARS-CoV-2 has accrued mutations throughout its 30kb genome. Of particular interest are the mutations present in the ORF8 protein, which occur in every major variant. The precise function and impact of this protein on disease severity and pathogenesis remains understudies. Our studies reveal that the ORF8 protein modulates the immune response by impacting macrophage infiltration into the lungs. Additionally, we have shown that the ORF8 protein of SARS-CoV-2 has accrued mutations throughout its evolution that lead to a loss of function phenotype in this protein. Our work reveals that the ORF8 protein of SARS-CoV-2 contributes significantly to disease progression through modulation of the inflammatory response.

## Introduction

A cluster of viral pneumonia cases was first observed in Wuhan, Hubei province, China in December of 2019. The causative agent of this pneumonia was later revealed to be a novel coronavirus, now known as severe acute respiratory syndrome coronavirus 2, or SARS-CoV-2. The virus quickly spread, leading to the declaration of a pandemic in 2020.^1^ The pandemic, known as the COVID-19 pandemic, has now claimed over 6.9 million lives as of September 2023.^2^ Despite the widespread deployment of vaccines, the pandemic persists, with the emergence of viral variants greatly affecting vaccine effectiveness.

SARS-CoV-2 is a beta coronavirus that shares significant homology to SARS-CoV-1, which was responsible for viral pneumonia cases originating in China in 2002.^3,4^ The enveloped virus possesses a 30kb positive-sense RNA genome. The genome is functionally divided into thirds, with the 5′ end of the genome encoding the replication machinery and the 3′ end encoding the structural proteins Spike (S), Membrane (M), Envelope (E), and Nucleocapsid (N). Interspersed with the structural proteins are a variety of accessory proteins.^5^ All coronaviruses possess accessory proteins, although the number of accessory proteins and their functions vary amongst the members of the coronavirus family. The accessory proteins of coronaviruses do follow a functional theme, with many of the proteins being implicated in antagonism of both Type I Interferon (IFN) signaling and activity of Interferon-Stimulated Genes (ISGs), and others being implicated in interference with the cellular autophagy process which it uses to acquire membrane sources for viral replication.^6^

Many of the accessory proteins of SARS-CoV-2 possess high sequence homology, of around 80%, to accessory proteins found in SARS-CoV-1.^7^ The ORF8 protein of SARS-CoV-2 is the most divergent accessory protein from SARS-CoV-1, with the proteins only possessing 40% amino acid identity.^4^ A main difference between the two is that the ORF8 of SARS-CoV-1 possesses a 29 base pair deletion that divides the ORF8 into two separate ORFs, known as ORF8a and ORF8b.^4,6^ The function of SARS-CoV-1 ORF8a and ORF8b is hypothesized to involve the degradation of IRF3.^8^ SARS-CoV-2 ORF8 has numerous hypothesized functions, including down-regulation of MHC class I on the cell surface, modulation of spike incorporation into virions, and agonism of IL-17 receptor signaling.^9–11^

Our previous work with accessory protein deletion viruses of SARS-CoV-2 revealed that an ORF8 deletion virus resulted in increased lung inflammation when compared to a clinical isolate.^12^ To further characterize the impact of ORF8 in clinical disease progression, we infected mice with two different doses of our WA-1ΔORF8 virus and compared weight loss and lung inflammation to a clinical isolate of WA-1. Our results show that the absence of ORF8 impacts inflammation at both doses, causing increased immune cell infiltration. As every variant that has emerged since B.1.1.7 in late 2020 possesses either one or more ORF8 mutations, we aimed to characterize the impact of these naturally occurring ORF8 mutations on viral pathogenesis by synthesizing recombinant WA-1 virus containing variant ORF8 genes of B.1.1.7, B.1.351, and P.1 to characterize the functional consequences of these mutations in a murine model.

## Materials and Methods

### Infectious Clone Construction and Rescue

#### SARS-CoV-2 DNA Fragment Cloning

The cloning of SARS-CoV-2 DNA fragments and transformation-associated recombination (TAR) assembly of genomes were performed as previously described.^12^ Briefly, the SARS-CoV-2 WA1 genome was cloned into seven individual DNA fragments: 1a-1, 1a-2, 1a-3, 1b-1, 1b-2, S, and AP. The TAR vectors were PCR amplified from pCC1BAC-his3 and used to clone each fragment. Yeast transformants were patched on synthetic dropout medium plates, and correct junctions between the fragment and vector were confirmed by detection PCR and Sanger sequencing.^13^ Plasmid DNAs of the fragments were isolated from E. coli and used for complete genome assembly or further modifications.

#### ORF8 Deletion in AP Fragment

The generation of ORF8 deletions in the AP fragment was achieved using in vitro CRISPR-Cas9 digestion and TAR assembly, as described in our previous publication.^12^ The process involved Cas9 target site identification, sgRNA transcription, AP fragment digestion, and TAR assembly with an oligo in yeast via spheroplast transformation.

#### Generation of AP Fragments with ORF8 Single Mutations and ORF8 Genes in B.1.1.7 and P.1 Variants

To generate ORF8 single mutations (S84L, E92K) and ORF8 genes in B.1.1.7 and P.1 variants, short fragments were amplified using Platinum SuperFi II DNA polymerase (Thermo Fisher) and the wild-type AP fragment as a template. The ORF8 single mutations and ORF8 genes in B.1.1.7 and P.1 variants were introduced by primers into each amplicon, which had 30-35 bp of homology at each end between the adjacent fragments. The amplicons were then digested with DpnI (NEB) to remove template DNA and purified with a Qiagen PCR purification kit.

Subsequently, 50 fmol of each amplicon and 15 fmol of the YCpBAC vector were assembled using the standard Gibson assembly reaction (NEB), transformed into *E. coli* DH10B competent cells (Thermo Fisher), and plated on LB medium containing 12.5 mg/ml chloramphenicol. The presence of desired mutations or ORF8 genes in B.1.1.7 and P.1 variants was confirmed through colony PCR and Sanger sequencing (GeneWiz). Plasmid DNAs containing the desired modifications were isolated from *E. coli* using the Purelink HiPure Plasmid Midiprep Kit (Thermo Fisher).

#### Complete Genome Assembly

The complete genome assembly was performed as previously described.^12^ Briefly, the TAR vector for the complete genome was generated by assembling a pCC1-ura3 amplicon, a CMV promoter amplicon, a BamHI fragment, a polyA fragment, and an HDVR+BGH region into circular DNA in yeast by TAR (Table S4). SARS-CoV-2 DNA fragment plasmids were digested with I-SceI (NEB) to release the overlapping fragments from the backbones, and the complete genome vector was linearized by BamHI (NEB) digestion. The fragments and complete genome vector were mixed with yeast spheroplasts for TAR assembly. Transformants were patched on SD-URA plates, and positive clones were screened by PCR. The subsequent DNA isolation from yeast, transformation into, and extraction from *E. coli* followed the same procedure as for DNA fragment clones.

For a detailed description of the complete genome assembly, please refer to our previous publication^12^.

#### Virus Reconstitution

24 hours prior to transfection, 5e4 VeroTMPRSS2 cells (*ATCC*, Manassas, VA) were plated per well in 1mL of VeroTMPRSS2 media (DMEM (*Quality Biological,* Gaithersburg, MD), 10% FBS (*Gibco,* Waltham, MA), 1% Penicillin-Streptomycin (*Gemini Bio Products*, Sacramento, CA), 1% L-Glutamine (*Gibco,* Waltham, MA)). For transfection, 5μg of the infectious clone and 100ng of a SARS-CoV-2 WA-1 nucleoprotein expression plasmid were diluted in 100μL of OptiMEM (*Gibco,* Waltham, MA). 3μL of TransIT-2020 (*Mirus Bio,* Madison, WI) was added and the reactions were incubated for 30 minutes prior to addition to the cells in the BSL-3. Cells were checked for cytopathic effect (CPE) 72-96 hours after transfection and the supernatant collected for plaque purification. For plaque purification, 6e5 VeroTMPRSS2 cells were plated in a 6-well plate in 2mL of VeroTMPRSS2 media 24 hours prior to infection. 25μL of this supernatant was then serial diluted 1:10 in DMEM and 200μL of this supernatant was added to the VeroTMPRSS2 cells. The cells were rocked every 15 minutes for 1 hour at 37°C prior to overlay with 2mL of a solid agarose overlay (EMEM (*Quality Biological,* Gaithersburg, MD), 10% FBS, 1% Penicillin-Streptomycin, 1% L-Glutamine, 0.4% w/v SeaKem agarose (*Lonza Biosciences,* Morrisville, NC). Cells were incubated for 72 hours at 37°C and 5% CO_2_ and individual plaques were picked and then transferred to a well of a 6-well plate with 4e5 VeroTMPRSS2 cells in 3mL VeroTMPRSS2 media. After 48 hours, successful plaque picks were assessed by presence of CPE. 1mL of a well showing CPE was transferred to a T175 with 8e6 VeroTMPRSS2 cells in 30mL of VeroTMPRSS2 media and the virus stock was collected 48 and 72 hours after. The stocks were then titered by plaque assay.

### Titering of Virus Stocks, Growth Curve Samples, Tissue Homogenates by Plaque Assay

The day prior to infection, 2e5 VeroTMPRSS2 cells were seeded per well in a 12-well plate in 1mL of VeroTMPRSS2 media. Tissue samples were thawed and homogenized with 1mm beads in an Omni Bead ruptor (*Omni International Inc.*, Kennesaw, GA) and then spun down at 21,000xg for 2 minutes. A 6-point dilution curve was prepared by serial diluting 25μL of sample 1:10 in 225μL DMEM. 200μL of each dilution was then added to the cells and the plates were rocked every 15 minutes for 1 hour at 37°C. After 1hr, 2mL of a semi-solid agarose overlay was added to each well (DMEM, 4%FBS, 0.06% UltraPure agarose (*Invitrogen*, Carlsbad, CA). After 72 hours at 37°C and 5% CO_2_, plates were fixed in 2% PFA for 20 minutes, stained with 0.5mL of 0.05% Crystal Violet and 20% EtOH, and washed 2x with H_2_O prior to counting of plaques. The titer was then calculated. For tissue homogenates, this titer was multiplied by 40 based on the average tissue sample weight being 25mg.

### Growth Curve Infection and Sample Processing

The day prior to infection, 1.5e5 VeroTMPRSS2 cells or 1.5e5 A549-ACE2 cells (Courtesy of Dr. Brad Rosenberg, *Icahn School of Medicine at Mount Sinai)* were seeded in a 12-well plate in 1.5mL of VeroTMPRSS2 media or A549-ACE2 media (DMEM, 10% FBS, 1% penicillin-streptomycin). The day of infection, cells were washed with 500μL of DMEM and the volume of virus needed for an M.O.I. of 0.01 was diluted in 100μL DMEM and added to a well in triplicate. The plates were rocked every 15 minutes for 1hr at 37°C. After 1hr, the inoculum was removed, the cells were washed with 500μL of complete media and then 1.5mL of complete media was added to each well. 300μL of this supernatant was taken as the 0hr timepoint and replaced with fresh media. 300μL of supernatant was pulled and replaced with fresh media at 0hr, 6hr, 24hr, 48hr, 72hr, and 96hr timepoints. This supernatant was then titered.

### Infection of hACE2-k18 Mice

All animals were cared for according to the standards set forth by the Institutional Animal Care and Use Committee at the University of Maryland, Baltimore. On Day 0, 12-14 week old hACE2 transgenic K18 mice (K18-hACE2) (*Jackson Labs,* Bar Harbor, ME) were anesthetized interperitoneally with 50μL ketamine (1.3mg/mouse)/xylazine (0.38mg/mouse). The K18-hACE2 mice were then inoculated with either 1e2 or 1e3 PFU of each virus in 50μL PBS. The mock infected mice received 50μL PBS only. Mice were then weighed every day until the end of the experiment. Mice were euthanized with isoflurane on day 2, 4 and 7. From each mouse, the left lung was collected in PFA for histology and the right lung was split in half with one half placed in PBS for titer and one half placed in TRIzol for RNA extraction.

### RT-qPCR of Tissue Homogenates

Samples were homogenized in an Omni Bead ruptor in TRIzol and then the RNA was extracted from 300μL of each sample using the Direct-zol RNA miniprep kit (*Zymo Research,* Irvine, CA). 2μL of isolated RNA from each sample was then converted to cDNA using the RevertAID first strand cDNA synthesis kit (*Thermo Fisher,* Waltham, MA) in a 20μL total reaction volume. For qPCR for SARS2 Rdrp, 20μL reactions were prepared using 2μL cDNA, 1μL of 10mM Rdrp Forward primer (10006860, *Integrated DNA Technologies*, Coralville, IA), 1μL of Rdrp Reverse primer (10006881, *Integrated DNA Technologies*, Coralville, IA), and 10μL of 2x SYBR Green (*Thermo Fisher,* Waltham, MA). The reactions were then run on a 7500 Fast Dx Real-Time PCR Instrument (4357362R, *Applied Biosystems*, Waltham, MA). For qPCR for murine GAPDH, 20μL reactions were prepared using 2μL cDNA, 1μL of a 20x murine GAPDH primer (MM.pt.39a.1, *Integrated DNA Technologies*, Coralville, IA), and 10μL of 2x SYBR Green. The reactions were then run on a QuantStudio 5 Real-Time PCR Instrument (A28133, *Applied Biosystems*, Waltham, MA).

### H/E Staining of Lungs and Pathological Scoring

Lungs were scored in a blinded fashion on a scale from 0 to 5 with 0 being no inflammation and 5 being the highest degree of inflammation. Interstitial inflammation and peribronchiolar inflammation were scored separately. Scores were then averaged for the overall inflammation score.

### Cytokine Arrays

The concentration of lung RNA was quantified using a Nanodrop (NanoVue Plus, *GE Healthcare,* Chicago, IL). 400ng of RNA was converted to cDNA using the Qiagen RT^2^ First Strand Kit (330404, *Qiagen*, Hilden, Germany). The cDNA was analyzed with the Qiagen RT^2^ Mouse Cytokines and Chemokines array (PAMM-150Z, *Qiagen*, Hilden, Germany). Reactions were run on a QuantStudio 5 Real-Time PCR Instrument (A28133, *Applied Biosystems*, Waltham, MA). The results were analyzed with the Qiagen analysis spreadsheet provided with the kit.

### RNA Sequencing and Analysis

Library preparation and sequencing was performed by the University of Maryland Institute of Genome Sciences (Baltimore, MD, USA). After RNA extraction, as described above, transcriptomic libraries were generated and sequenced on an Illumina NovaSeq 6000 (S4 flow cell, 100bp paired-end; *Illumina*, San Diego, CA). Raw data is available in the NCBI SRA under the accession number PRJNA857920. Reads were preprocessed using cutadapt v3.4, then aligned to the murine genome (assembly GRCm38) using STAR v2.7.8a.^14,15^ Genes with at least a mean count of 10 reads in at least one condition were subject to differential expression analysis with DESeq2 v4.1.0 followed by pathway analysis using Ingenuity Pathway Analysis (*Qiagen*, Hilden, Germany).^16^ Genes were only considered for follow-up if the magnitude of differential expression was at least 2-fold in either direction and the difference in expression between conditions was significant (p<0.05) after multiple testing correction.

### Immunostaining for Immune Cell Markers

Histological samples were processed according to optimized protocols established by *HistoWiz* (Brooklyn, NY, USA). Samples were stained using antibodies for SARS-CoV-2 nucleocapsid, the macrophage marker F4-80 (*Cat #14-4801-82, Clone BM8*), and the neutrophil marker Ly6G (*Cat #ab25377, Clone RB6-8C5*). Images were taken using the *HistoWiz* software.

### Statistical Analysis

All statistical analyses were carried out using the GraphPad Prism software (GraphPad Software, San Diego, CA, USA) or R version 4.1.1.^17^ The cutoff value used to determine significance was p≤0.05 for all tests. The statistical tests run were unpaired t-tests assuming unequal variances, or one-way ANOVA followed by T-test with Bonferroni correction, where indicated. For the differential expression analysis, reported p-values are the result of the Wald test after Benjamini-Hochberg correction, as calculated by DESeq2 v4.1.0.^16^

### Biosafety approval

All virus experiments and recombinant virus creation was approved by the Institutional Biosafety Committee at The University of Maryland, Baltimore.

## Results

### Infection of K18-hACE2 mice with WA-1ΔORF8 results in increased weight loss and increased lung inflammation compared to mice infected with WA-1

SARS-CoV-2 ORF8 has been proposed to have several functions. To directly assess the role of ORF8 in pathogenesis we produced a recombinant SARS-CoV-2 virus lacking the complete sequence of ORF8 and compared pathogenesis of this mutant to the wildtype SARS-CoV-2 strain WA-1. Our previous work with an ORF8 deletion virus revealed a difference in lung inflammation when compared to mice infected with the wildtype virus. To further investigate this phenotype, we infected 12-14 week old K18-hACE2 transgenic mice with either 1e2 PFU or 1e3 PFU of either WA-1 or WA-1ΔORF8. We assessed weight loss daily, lung viral loads by plaque assay, and viral RNA levels by qPCR for RdRp were measured at days 2, 4, and 7 post-infection. At both doses, mice infected with WA-1ΔORF8 lost weight a day earlier than mice infected with the full-length WA-1 (**Figure 1A, B**). By day 7, mice infected with the ORF8 deletion virus had 5-7% more weight loss compared to mice infected with the full-length virus (**Figure 1A, B**). Despite the increased weight loss, no differences in lung viral titer were observed at the 1e2 PFU dose (**Figure 1C**). In the 1e3 PFU dose, we find that WA-1ΔORF8 infected mice have a statistically significant one log decrease in lung viral titer at Day 4 compared to the WA-1 infected mice (p=0.012, **Figure 1C**). Levels of viral RNA by qPCR for RdRp in the WA-1ΔORF8 mice were measured and no statistically significant changes were observed at both doses across all days (**Figure 1D**).

**Figure 1.**
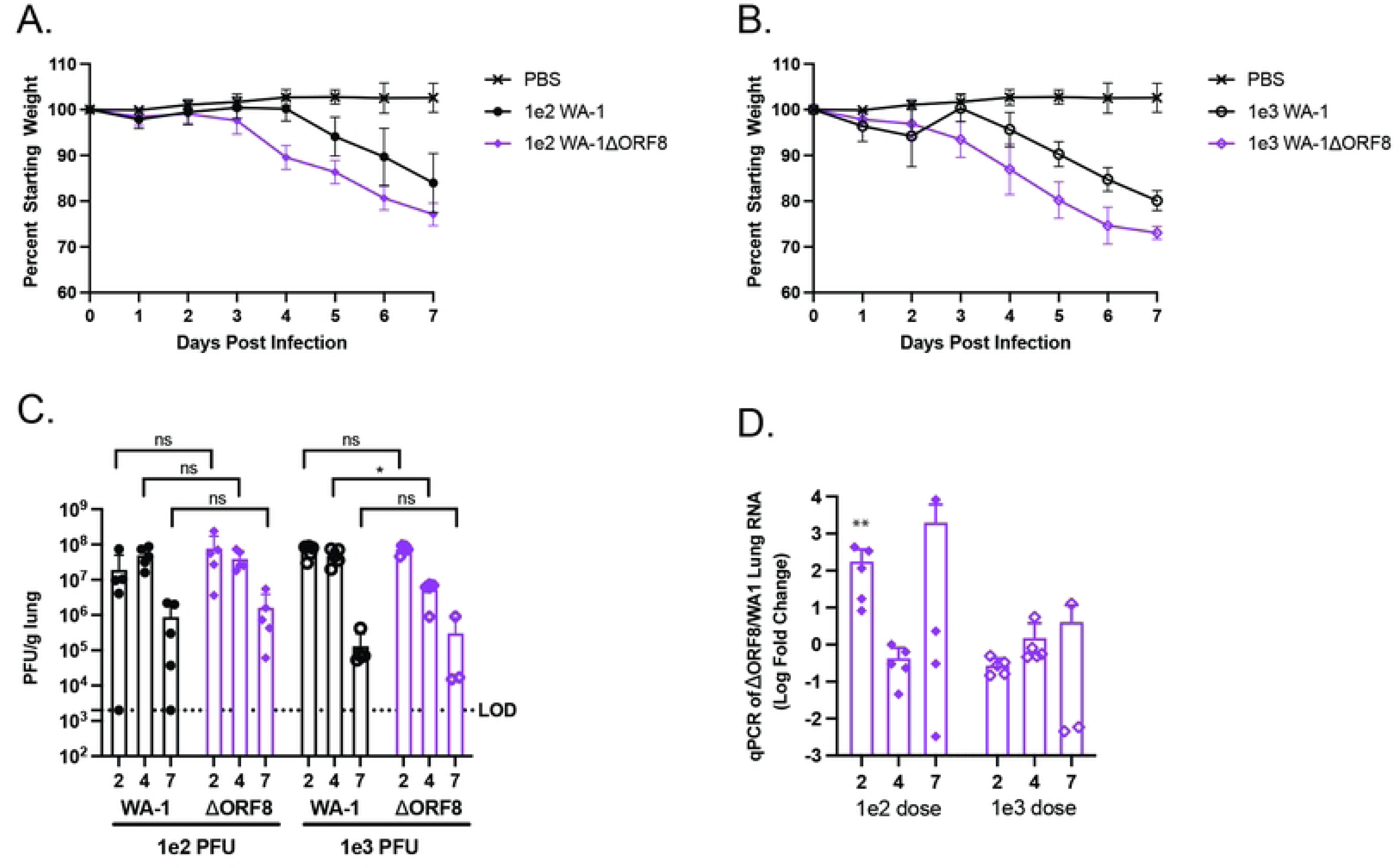
Pathogenesis of K18/hACE2 mice infected with WA1 or WA-1ΔORF8. K18/hACE2 mice were infected with either WA1 or WA-1ΔORF8 to determine differences in pathogenesis. A. Weight loss of mice infected with either PBS, 1e2 PFU of SARS-CoV-2 WA-1, or 1e2 PFU of SARS-CoV-2 WA-1ΔORF8 (n=15 mice per virus). B. Weight loss of mice infected with either PBS, 1e3 PFU of SARS-CoV-2 WA-1, or 1e3 PFU of SARS-CoV-2 WA-1ΔORF8. (n=15 mice per virus) C. Live virus titers by PFU/g lung in mice infected with either WA-1 or WA-1ΔORF8. (n=5 mice per timepoint) D. Viral RNA levels of RdRp expressed as a fold change relative to the comparable dose of WA-1 (n=5 mice per timepoint). Sample comparisons with significant differences are shown. (*, p≤ 0.05; **, p≤0.005; ***, p≤0.0005, ns=not significant).

The lungs from these mice were fixed and stained for histological analysis. Despite the fact that mice infected with 1e2 WA-1ΔORF8 lost more weight than mice infected with 1e2 WA-1, the lungs exhibited similar levels of inflammation (**Figure 2A,B**). At the dose of 1e3 PFU, mice infected with the deletion virus exhibited increased inflammation in the lungs at days 2, 4, and 7 post-infection. This inflammation included infiltration of cells into the alveolar space and both thickening and sloughing of cells of the major airways (**Figure 2A**). Histological scoring of the lungs revealed statistically significant differences, with the lungs of the mice infected with 1e3 WA-1ΔORF8 exhibiting increased inflammation at days 2, 4, and 7 post-infection (Day 2 p=0.027, Day 4 p=0.0013, Day 7 p=0.0027).

**Figure 2.**
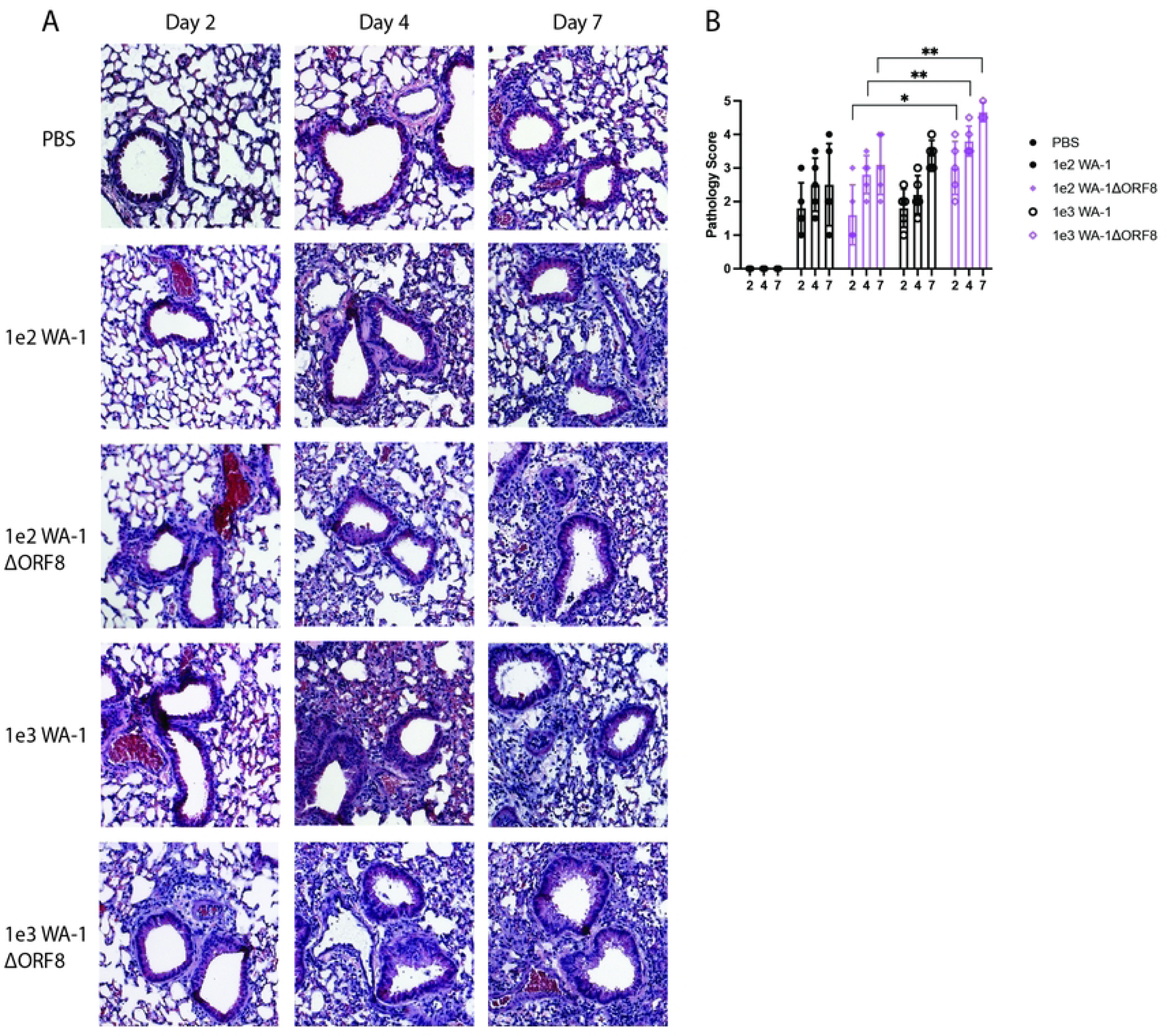
Lung histology and histological scores of K18/hACE2 mice infected with WA-1 or WA-1ΔORF8. Lungs from infected mice at day 2, 4 and 7 post infection were fixed and sectioned for staining with H&E. A. H&E staining of the lungs of mice infected with WA-1 or WA-1ΔORF8. Representative images of each group, n=5 mice per virus and timepoint. B. Histological scoring of the lungs of mice infected with either WA-1ΔORF8 or WA-1. Scoring described in methods section (n=5 mice per timepoint) Sample comparisons with significant differences are shown. (*, p≤ 0.05; **, p≤0.005; ***, p≤0.0005).

### Transcriptomic analysis of the lungs of K18-hACE2 mice infected with WA-1ΔORF8 or WA-1 reveals an upregulation of macrophage signaling pathways and a concurrent increase in the population of macrophages in the lungs by immunohistochemistry

To investigate the pathways associated with the increased inflammation seen in the lungs of mice infected with the WA-1ΔORF8 virus, we performed an analysis of the inflammatory cytokines and chemokines in the lungs of these mice at Days 2, 4 and 7, finding that many cytokines and chemokines associated with innate immune cell chemotaxis and cytokine storm signaling were upregulated early in infection (**Table 1**). Notably, we observed more significant changes in the lower dose (1e2 PFU) as compared to the higher dose (1e3 PFU), which we hypothesize is due to the increased amount of virus masking any subtler changes.

**Table 1.**
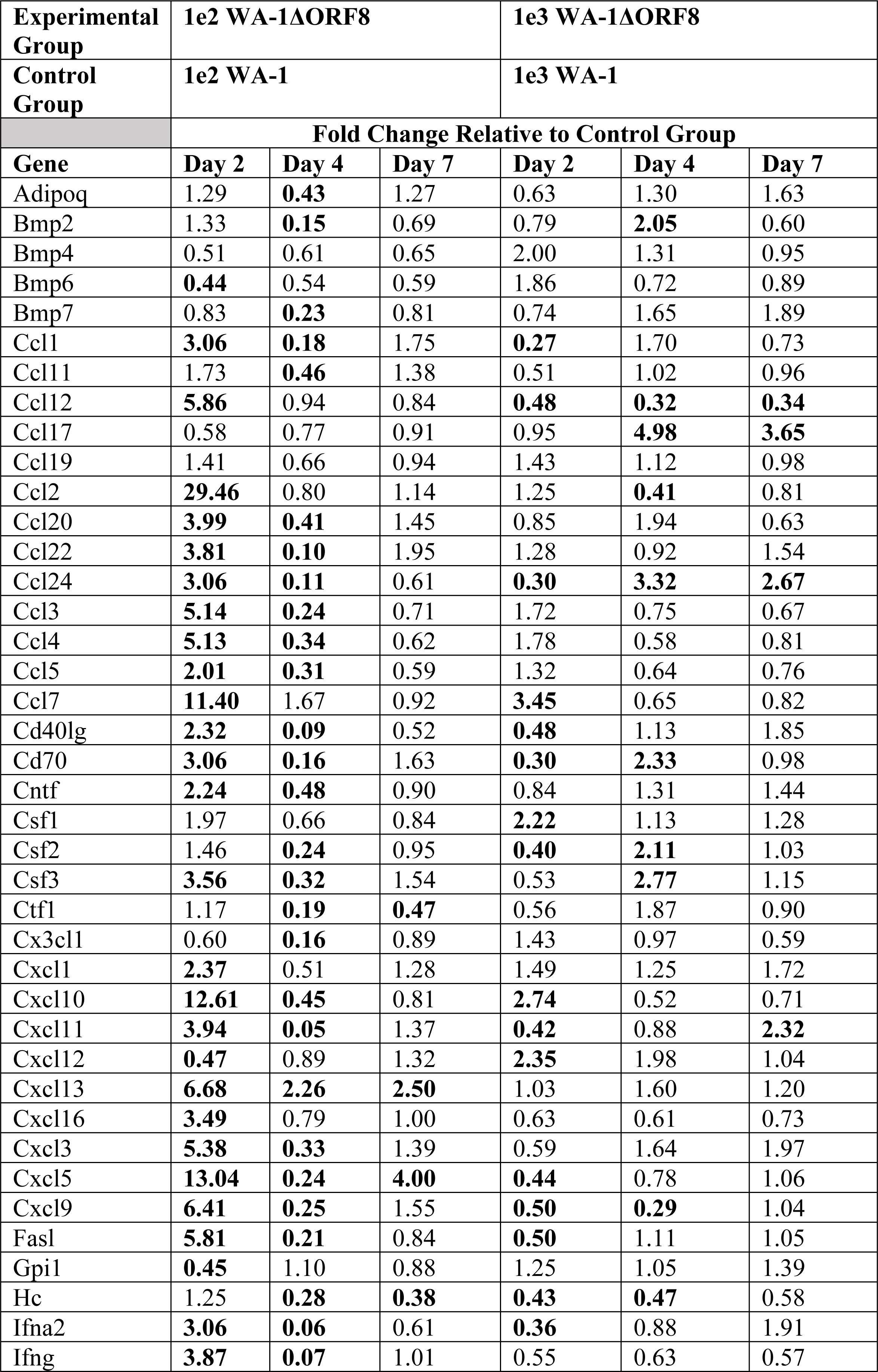
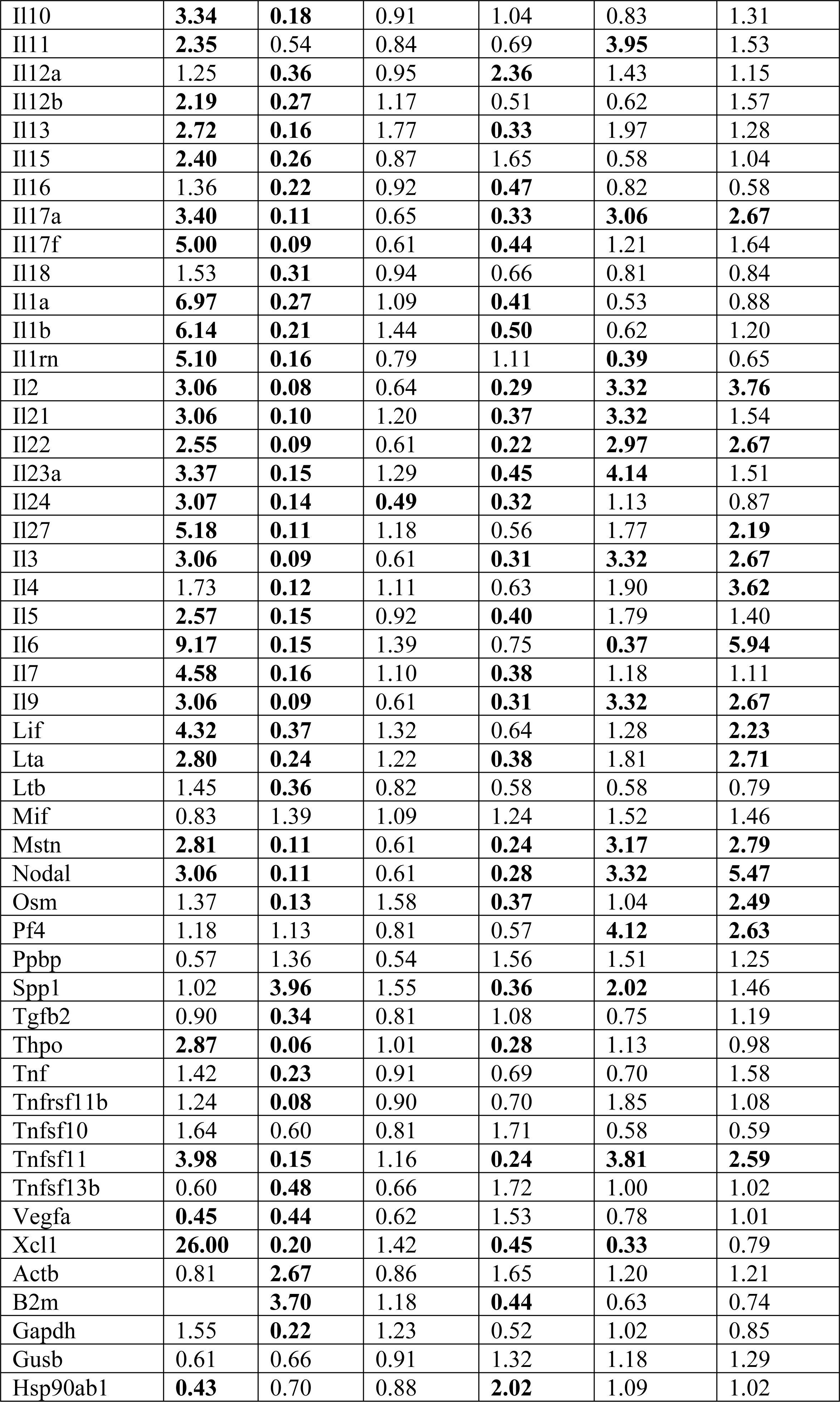
Fold changes of inflammatory cytokines and chemokines of mouse lungs on Day 2, Day 4, and Day 7. The values in bold are statistically significant.

Specifically, we noticed an induction of key inflammatory proteins that recruit or stimulate macrophages and monocytes in the lungs like CCL1, CCL2, CCL4, CCL7 and CXCL10 (**Table 1**). This demonstrates an increase in chemokines involved with recruiting macrophages and neutrophils, which correlates with what is observed in the H&E scoring in **Figure 2**. For a further analysis of the affected pathways, we isolated the RNA from the lungs of mice infected with either the deletion virus or the full-length virus and performed RNA-seq. Transcriptomic analysis of mouse lungs infected with WA-1 or WA-1ΔORF8 at days 2, 4 and 7 post infection identified key changes to the immune response in the lungs dependent on the virus (**Figure 3A-C**). In particular, neutrophil associated responses were similar across time points for both viruses. Classical macrophage (M1 macrophages) associated genes were differentially regulated early in infection at day 2. WA-1 infected lungs have a muted induction of M1 macrophage associated genes while WA-1ΔORF8 infected mice have significant and rapid induction of M1 macrophage associated genes. This includes CD68, CD80 and CD86 (**Figure 3D**). M2 macrophage associated genes were also differentially upregulated in WA-1ΔORF8 infected mice with ARG1, CLEC7a and RETNLB significantly upregulated compared to WA-1 infected mice (**Figure 3E**). Comparison of fold change of WA-1ΔORF8 / WA-1 infected mice for these selected genes demonstrates early induction of the macrophage/monocyte induced genes in the WA-1ΔORF8 infected mice (**Figure 3F and G**). This suggests that the inflammation observed in lungs is a result of macrophage activation and recruitment that may be impairing lung function to cause more severe disease, as we observe with the WA-1ΔORF8 virus.

**Figure 3.**
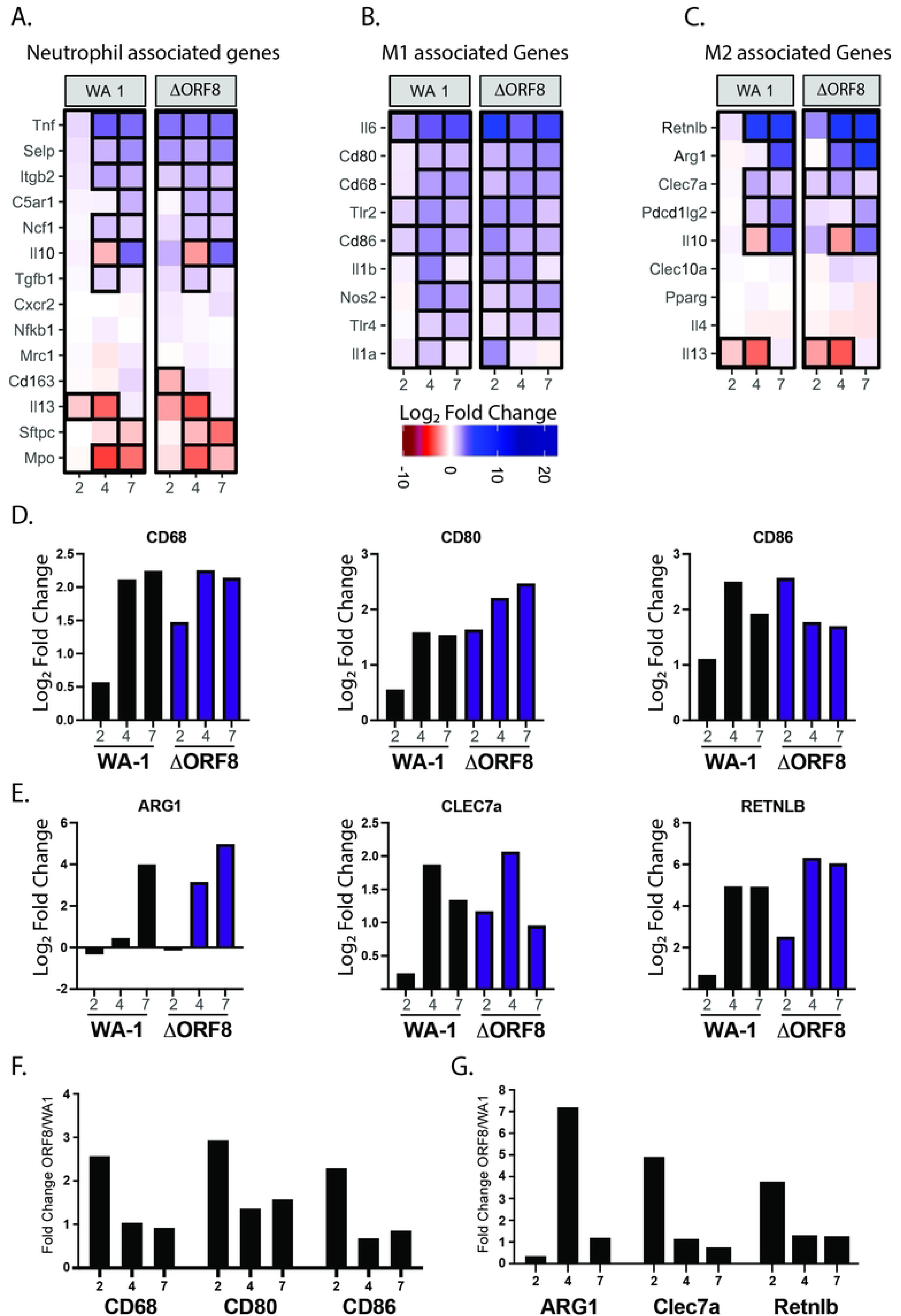
Differentially expressed genes in WA-1 and WA-1ΔORF8 infection. A, B and C. Selected genes associated with neutrophils (A), M1 macrophages (B) and M2 macrophages C) are shown as heat maps. Differentially expressed pathways at Day 2, 4 and 7. All Log_2_ fold change is relative to a PBS control infection mouse at the same timepoint. A scale of fold change is shown below. D. Graphing of Log_2_ fold change for select genes in each set in B and C that show dynamic changes in heat map. F and G. Fold change graphed from transcriptomic data of ΔORF8 compared to WA1 infections at each time point for selected genes.

Since we observed an increase in neutrophil and macrophage signaling pathways, we wanted to determine the localization of these cells in the lungs, particularly at the dose of 1e3 PFU at day 7 post-infection where we saw the most significant differences in lung inflammation between the mice infected with the deletion virus and the mice infected with the full-length virus. We performed immunohistochemistry on the lungs of these mice, staining for the markers Ly6G to identify neutrophils and F4-80 to identify macrophages. At day 7 post-infection, mice infected with WA-1 and WA-1ΔORF8 had similar insignificant levels of Ly6G positive neutrophils in the lungs. However, the lungs of the mice infected with the WA-1ΔORF8 virus had significantly more macrophages in the lungs at this timepoint, which correlates with the upregulation of macrophage signaling pathways seen in the lungs of these mice early in infection by RNASeq analysis (**Figure 4**). There was no difference in localization of the virus by nucleocapsid staining at this timepoint, with the majority of the virus being cleared from the large airways and localizing to the alveolar space.

**Figure 4.**
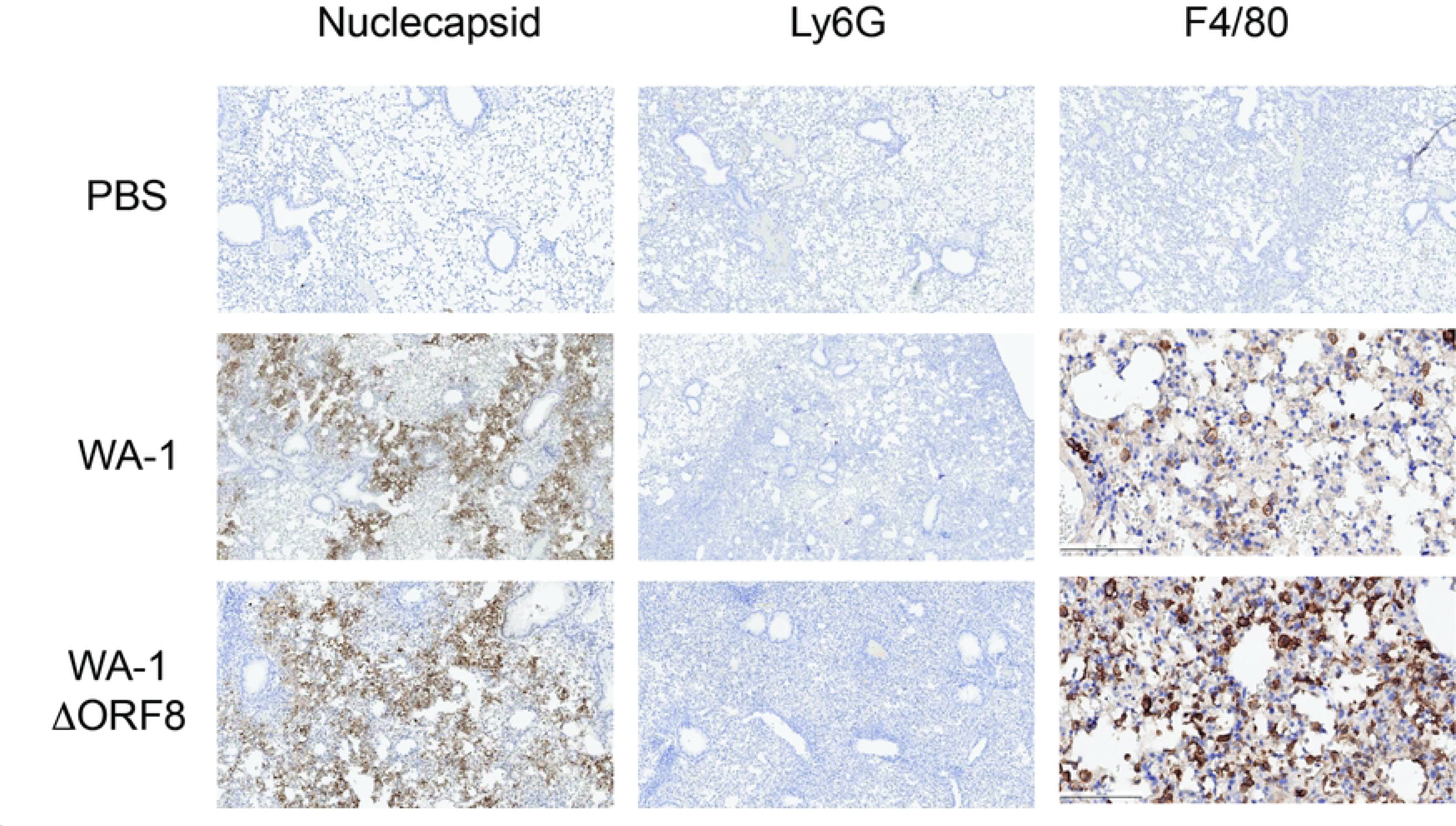
Staining for neutrophils (Ly6G), macrophages (F4-80) and nucleocapsid in the lungs of WA-1 and WA-1ΔORF8 infected mice. Lungs from WA-1 and WA-1ΔORF8 infected mice at 7-day post infection were fixed and stained for neutrophils (anti-Ly6G), macrophages (anti-F4/80) or SARS-CoV-2 nucleocapsid (anti-N). Brown stain denotes positive antibody labeling of respective protein targets. Representative images shown of 5 mice at day 7 post infection.

### Naturally occurring mutations in ORF8 result in increased levels of inflammation in the lungs

With the emergence of the variants of SARS-CoV-2, we noticed an increase in the incidence of naturally occurring mutations in the ORF8 protein. To determine if these naturally occurring mutations affect inflammation, we generated recombinant WA-1 viruses containing variant ORF8 genes. We produced four viruses to study including a WA-1 virus with the B.1.1.7 ORF8, a WA-1 virus with the P.1 ORF8, a virus with the point mutation S84L and a virus with the point mutation E92K. The B.1.1.7 ORF8 protein contains four mutations, with the most notable being the introduction of a premature stop codon at amino acid 27 out of 121. The point mutation S84L occurs in all variants of SARS-CoV-2, beginning with the emergence of B.1.1.7 in December of 2020. The P.1 ORF8 contains the S84L mutation and an additional E92K mutation, which we also synthesized individually.

After producing this panel of WA-1 viruses containing mutant ORF8 genes, we determined the growth of the viruses in VeroTMPRSS2 cells. All of the viruses grew comparably (**Figure 5A**). We next infected K18-hACE2 mice, first with the WA-1 virus containing the B.1.1.7 ORF8 and a virus containing the single mutation of ORF8 S84L. We compared their weight loss and lung titers to mice infected with either WA-1 or the WA-1ΔORF8 virus. Mice infected with the WA-1ΔORF8 virus and the B.1.1.7 ORF8 virus, lost weight a day earlier and by day 7 lost 5-7% more weight than the mice infected with WA-1. The mice infected with the WA-1 ORF8 S84L virus began losing weight the same day as the WA-1 mice, but by day 7 lost 5-7% more weight than the mice infected with WA-1, similar to the WA-1ΔORF8 mice (**Figure 5B**). As we saw in mice infected with the deletion virus, there were no significant differences in lung titer by plaque assay amongst any of the groups at all time points (**Figure 5C**). We then infected K18-hACE2 mice with the WA1 virus containing the P.1 ORF8 and a WA-1 virus containing only the ORF8 E92K mutation. The mice infected with the P.1 ORF8 containing virus exhibited an intermediate weight loss phenotype compared to the WA-1 and the WA-1ΔORF8 mice, while the mice infected with the ORF8 E92K containing virus exhibited a weight loss phenotype that more closely resembled that seen in WA-1 infected mice (**Figure 5D**). As with the previous set of ORF8 mutant WA-1 viruses, we did not see significant differences in lung titers by plaque assay amongst all of the infected mice (**Figure 5E**).

**Figure 5.**
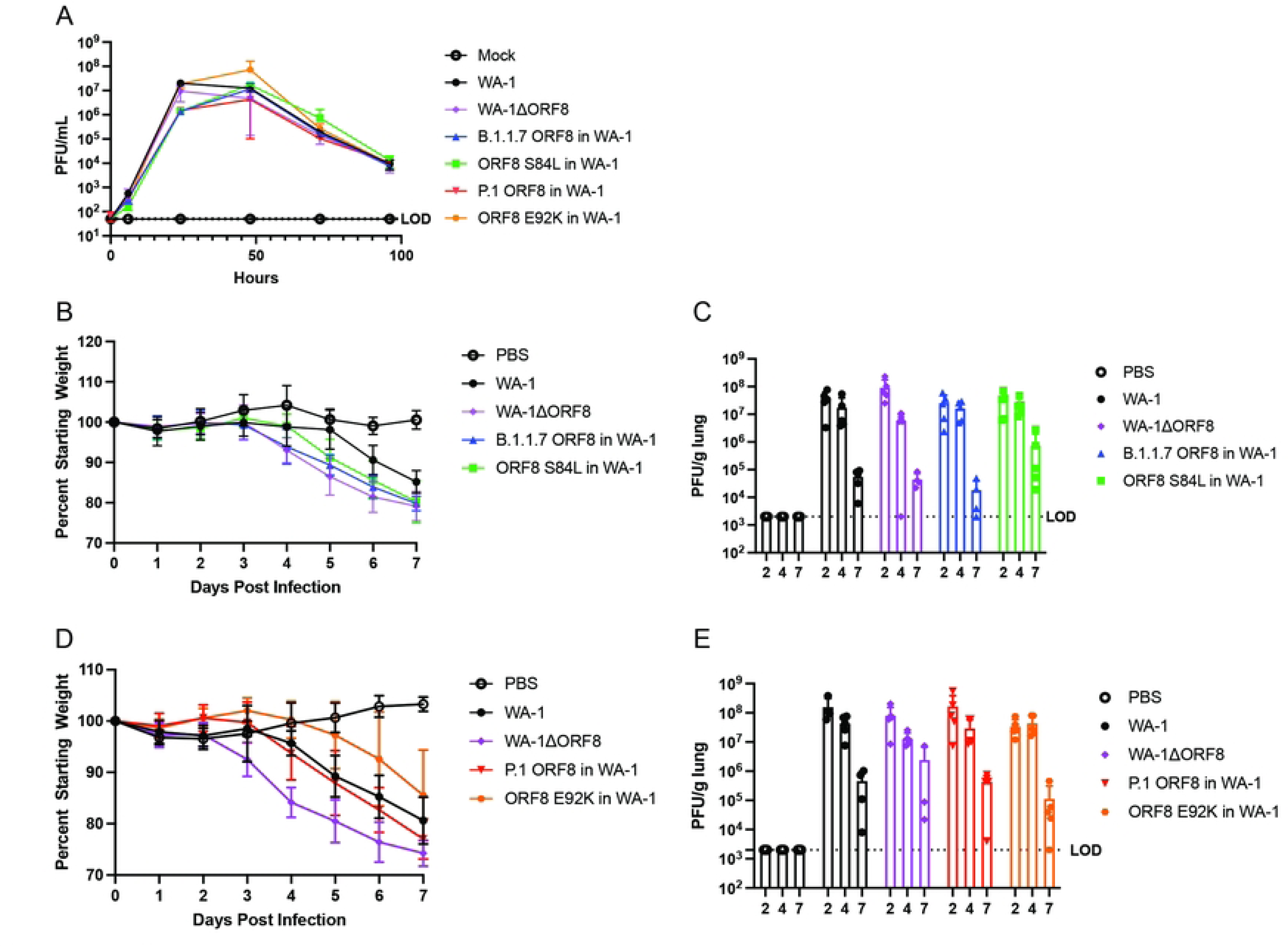
Infection of K18-hACE2 mice with variant ORF8 in WA-1 viruses. Cells and K18/hACE2 mice were infected with either WA1, WA-1ΔORF8, B.1.1.7 ORF8 in WA-1, or ORF8 S84L in WA-1 to determine differences in replication and pathogenesis. A. SARS-CoV-2 viruses expressing variant ORF8 genes were evaluated for growth by infection of VeroTMPRSS2 cells at MOI of 0.01 across a timecourse of infection. Supernatants were titered by plaque assay on VeroTMPRSS2 cells. B. Weight loss of K18-hACE2 mice infected with WA-1, WA-1ΔORF8, B.1.1.7 ORF8 in WA-1, or ORF8 S84L in WA-1. (n=5 mice/timepoint) C. Lung titers by plaque assay of K18-hACE2 mice infected with WA-1, WA-1ΔORF8, B.1.1.7 ORF8 in WA-1, or ORF8 S84L in WA-1. D. Weight loss of K18-hACE2 mice infected with WA-1, WA-1ΔORF8, P.1 ORF8 in WA-1, or ORF8 E92K in WA-1. E. Lung titers by plaque assay of K18-hACE2 mice infected with WA-1, WA-1ΔORF8, P.1 ORF8 in WA-1, or ORF8 E92K in WA-1.

We next examined the histopathological data of the lungs of mice infected with the panel of viruses. We noted significant inflammation in the lungs of all infected mice. However, as expected, the mice infected with the virus containing the B.1.1.7 ORF8, which contains a severely truncated ORF8, results in inflammation that is higher than the wildtype virus infected mice. These lungs contain a significant number of cells in the alveolar space and exhibit increased thickening of the major airways and sloughing of the epithelial cells into the air space (**Figure 6A**). Lung inflammation was scored for each virus (**Figure 6B**). Intriguingly, the lungs of mice infected with the ORF8 S84L in WA-1 virus also resemble the lungs of mice infected with the deletion virus, with a skew towards more inflammation in the lungs (**Figure 6A**).

**Figure 6.**
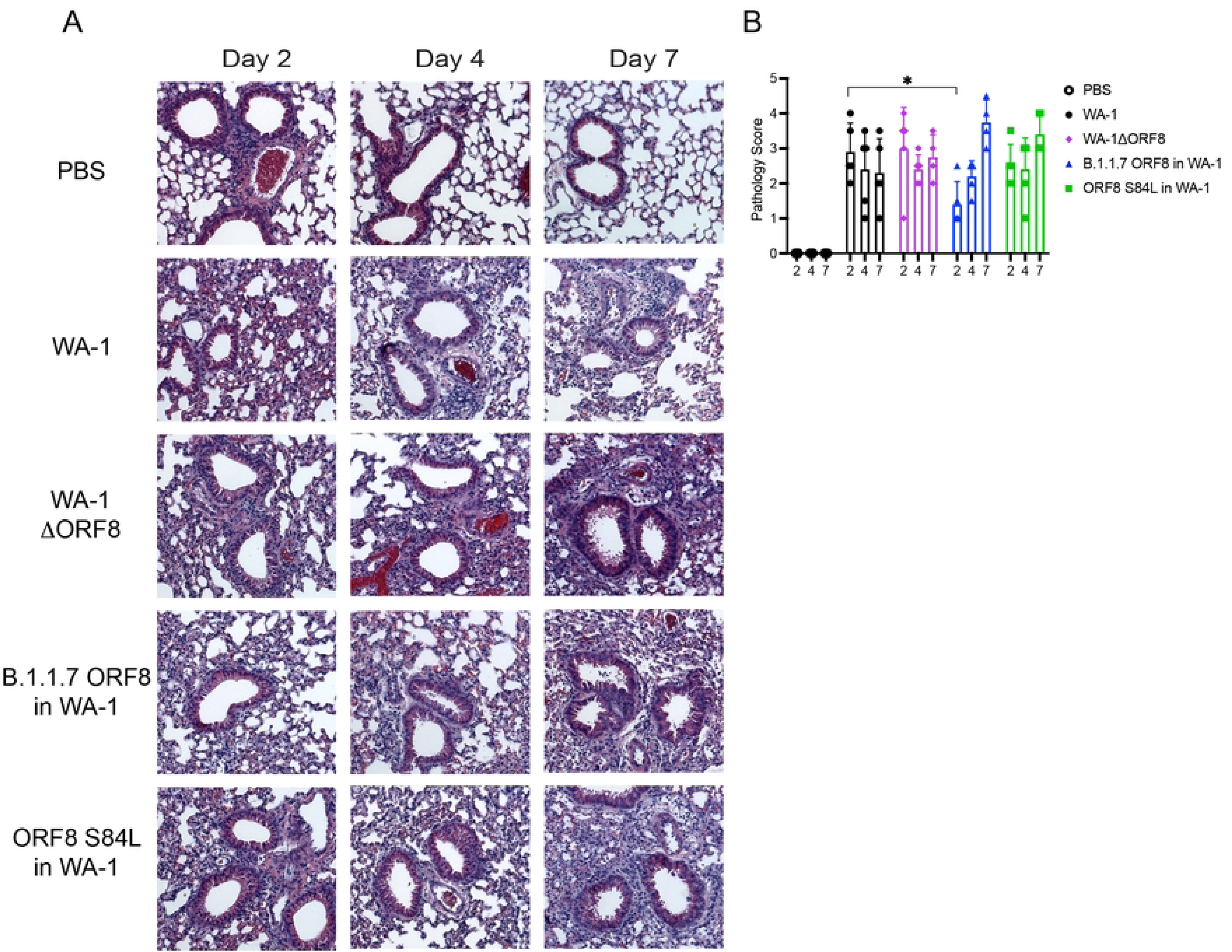
Lung histology and histological scores of mice infected with WA-1, WA-1ΔORF8, B. 1.1.7 ORF8 in WA-1, or ORF8 S84L in WA-1. Lungs from infected mice at day 2, 4 and 7 post infection were fixed and sectioned for staining with H&E. A. H&E staining of the lungs of mice infected with WA-1, WA-1ΔORF8, B. 1.1.7 ORF8 in WA-1, or ORF8 S84L. Representative images shown of 5 mice per virus per timepoint. B. Histological scoring of the lungs of mice infected with either WA-1ΔORF8 or WA-1. Scoring described in methods section (n=5 mice per timepoint). Sample comparisons with significant differences are shown. (*, p≤ 0.05; **, p≤0.005; ***, p≤0.0005).

The lungs of the mice infected with the P.1 ORF8 containing virus and the ORF8 E92K containing virus scored consistently with the weight loss data seen in **Figure 7**. The lungs of the mice infected with the P.1 ORF8 virus exhibited an intermediate inflammatory phenotype compared to the mice infected with WA-1 and the mice infected with the WA-1ΔORF8 virus. We observe more inflammation in the alveolar space when compared to WA-1, but less thickening of the major airways compared to what is observed in the mice infected with the ORF8 deletion virus (**Figure 7A**). The lungs of the mice infected with the deletion virus were more inflamed than the lungs of mice infected with the wildtype virus, with a striking significantly higher pathology score being seen at Day 7 (p=0.00016; **Figure 7B**). The P.1 ORF8 virus infected mice also had significantly higher inflammation in the lungs at Day 7 (p=0.0046) (**Figure 7B**). The lungs of the mice infected with the ORF8 E92K virus resembled the lungs of the mice infected with the WA-1 virus and did not achieve the levels of inflammation seen with the deletion virus and the P.1 ORF8 virus. However, at day 7, the lungs of these mice were more inflamed than the lungs of the mice infected with the wildtype virus (p=0.0061; **Figure 7B**).

**Figure 7.**
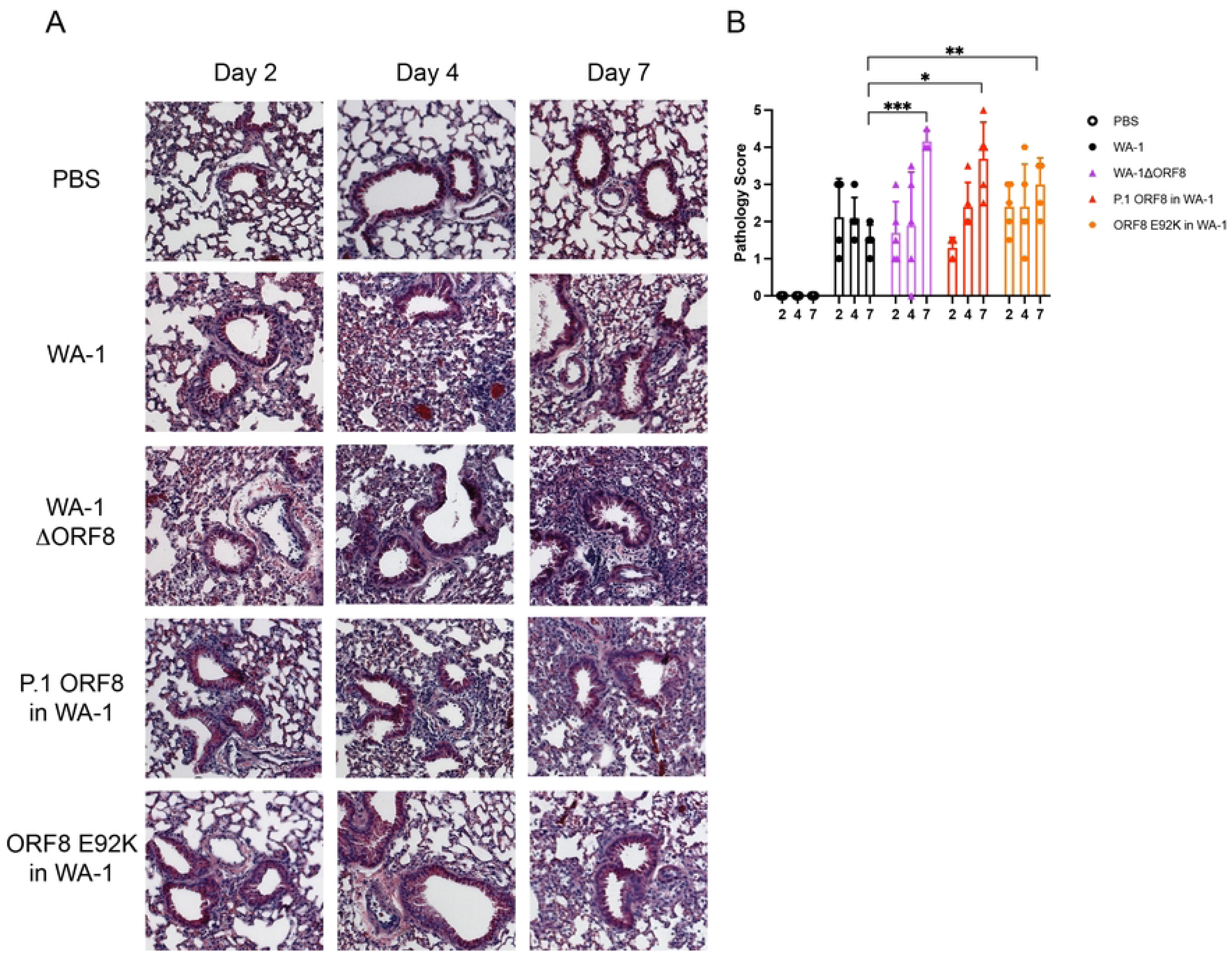
Lung histology and histological scores of mice infected with WA-1, WA-1ΔORF8, P.1 ORF8 in WA-1, or ORF8 E92K in WA-1. Lungs from infected mice at day 2, 4 and 7 post infection were fixed and sectioned for staining with H&E. A. H&E staining of the lungs of mice infected with WA-1, WA-1ΔORF8, P.1 ORF8 in WA-1, or ORF8 E92K. Representative images shown of 5 mice per virus per timepoint. B. Histological scoring of the lungs of mice infected with WA-1, WA-1ΔORF8, P.1 ORF8 in WA-1, or ORF8 E92K. Scoring described in methods section (n=5 mice per timepoint). Sample comparisons with significant differences are shown. (*, p≤ 0.05; **, p≤0.005; ***, p≤0.0005).

## Discussion

Our work with both an ORF8 deletion virus and variant ORF8 viruses has revealed that this protein contributes to modulating the inflammation caused by SARS-CoV-2. Mice infected with the WA-1ΔORF8 virus showed significantly higher levels of inflammation in the lungs compared to mice infected with the wildtype virus. Our bulk RNA sequencing data from the lungs of these mice revealed differential expression of genes involved in cytokine storm signaling pathways, macrophage activation pathways, and neutrophil signaling. In line with this RNA sequencing data, we saw a significant increase in the population of macrophages in the lungs of mice infected with the deletion virus compared to mice infected with the wildtype virus. We did not see a difference in the number of neutrophils in these lungs, suggesting that this is a cell-type specific phenomenon.

Previously published data suggests that the ORF8 of SARS-CoV-2 serves to downregulate MHCI, which presents antigen to CD8+ cytotoxic T cells.^9^ Our cytokine array data did show an increase in the transcript for β2-microglobulin (β2m), which would corroborate this data. However, we did not see MHCI appear in our bulk RNA sequencing data. This could mean that the downregulation of MHCI is mostly at the protein level and not at the transcript level. We did see a significant upregulation in the amount of macrophages in the lungs of the deletion virus-infected mice compared to the wildtype-infected mice. This may be due to signaling by the cytotoxic T cells, which secrete IFN-γ to activate macrophages.^18^ Notably, both T cells and macrophages are known to play a significant role in the immune response of SARS-CoV-2.^19,20^

Our work with the recombinant variant ORF8 viruses suggests that naturally occurring mutations in the SARS-CoV-2 ORF8 protein affect its function. As expected, the B.1.1.7 ORF8 in WA-1 mice exhibited increased weight loss and moderately increased inflammation in the lungs, which is unsurprising given that the ORF8 protein of the alpha lineage possesses a premature stop codon that truncates the protein 94 amino acids early. We have also shown that the mutation S84L, which is present in all of the variants, may serve to attenuate the function of ORF8, as mice infected with this virus and the P.1 ORF8 virus, which contains this mutation and an additional E92K mutation, appear to exhibit increased inflammation in the lungs compared to that caused by infection with the wildtype WA-1 virus. Current Omicron variants starting with the XBB lineage also have mutated ORF8 with a stop codon at amino acid 8 (G8*) in the ORF^1^. This leads to a truncated protein, similar to the mutation seen in B.1.1.7 where ORF8 is truncated at amino acid 27. Based on our data, the truncation in all XBB lineages should also lead to increased inflammation, which can result in beneficial or detrimental phenotypes in humans depending on the other mutations in the genome. Understanding how the individual mutations function by themselves as well as when combined with other mutations in the SARS-CoV-2 genome can help to predict the pathogenesis of future variants.

The predominant ORF8 mutation S84L arose early in the virus’ circulation. It is interesting to note that there were two predominant circulating strains of the original SARS-CoV-2 virus in Wuhan, China.^21,22^ These two strains were known as the S strain and the L strain, which were named for the amino acid they coded in the ORF8 protein. Interestingly, the L strain was attributed to more disease severity, which is corroborated by our data here. Given that the S strain gave rise to the L strain and the L strain is associated with more severe disease, it appears that there is some selective pressure on this ORF8 mutation that led to loss of function. It should be noted that the accessory protein ORF8 is the most highly divergent accessory ORF when compared to the ORF8 proteins of SARS-CoV-1. The SARS-CoV-1 ORF8 possesses a 29-nucleotide deletion that functionally splits the ORF8 region into two smaller proteins, ORF8a and ORF8b.^23^ Given our work and other published data, it appears that the loss of function of the ORF8 protein of SARS-CoV-2 has occurred independently throughout the evolution of the virus, suggesting that this loss of function is advantageous to the virus.^24–27^ Further studies aim to directly compare the S and L strains of Wuhan-1, characterize the differences in inflammation seen with the variant ORF8 proteins, and further investigate naturally occurring truncations and deletions in ORF8 and their effects on viral pathogenesis.

## Acknowledgements

This work was supported by grants from The Bill and Melinda Gates Foundation INV-006099, and DHS/BARDA ASPR-20-01495 to MF. Funding to SV was provided by NIH R01AI137365, NIH R03AI146632 and the J Craig Venter Institute.

